# Phylogenies as graphs: structured neural networks improve host origin predictions from paramyxovirus sequences

**DOI:** 10.64898/2026.09.18.752634

**Authors:** James C Herzig, Haley Stone, Liam Brierley

## Abstract

Accurately identifying the host of a virus from its genome sequence is a task with important applications in zoonotic disease surveillance and filling data gaps for metagenomic sampling. Machine learning approaches have seen broad application in making host predictions directly from viral genome sequences. However, most host prediction models do not incorporate information on viral phylogeny, which is strongly correlated with both genome composition and host. We apply a novel graph neural network (GNN) approach which explicitly represents viral phylogeny in model architecture to predict hosts of origin within the paramyxoviruses. We conduct rigorous benchmarking against non-structured neural networks and predictions made using phylogeny alone, showing that GNNs carry distinct advantages over other methods when making predictions where training data is sparse. Validation across different phylogenetic scales shows that simple phylogenetic prediction is effective in many applications and that phylogeny contributes a large proportion of the predictive power of host prediction models, with genome compositional features providing additional power only for specific predictions outside the range of the training data. This novel modelling approach and model validation framework are flexible and can be applied to other viral families.

**Author summary:** Advances in genome sequencing technology in the last two decades have resulted in a huge increase in the number and diversity of available viral genome sequences. However, many of these genomes do not have reliable associated data regarding the host which the source virus infects. Predicting the host a virus infects based on its genome sequence is therefore important both for the surveillance of emerging diseases from animal reservoirs and to fill data gaps which will enable future research. Computational models using machine learning have been widely applied to this problem. However, most previous applications do not consider the phylogenetic relationships between viruses, which has a strong correlation with host. Here, we have applied graph neural networks to predict the hosts of the paramyxoviruses, a family of viruses that contains numerous endemic and emerging threats to human and animal health. Our approach allows us to explicitly represent the relationship between viral sequences in the model architecture, helping improve predictions. We thoroughly assess our model’s performance relative to alternatives, finding that phylogeny is very important to host prediction and that our novel graph neural network approach results in more accurate predictions compared to other methods.

## 1. Introduction

Metagenomic sequencing is now discovering viruses at an unprecedented rate, with more than 800,000 sequences deposited to NCBI Virus in 2025. This has greatly increased our understanding of the breadth of viral diversity present across host species and environments [1–4]. However, a large proportion of available sequences do not have reliable associated metadata such as host range [5], either due to environmental or waste water sampling in the absence of a host [6,7] or due to ambiguity in whether sample hosts are incidental spillover hosts or hosts that maintain transmission (sometimes termed ‘reservoirs’ in the context of zoonotic spillover, we term these ‘hosts of origin’). Meanwhile, zoonotic transmission remains a major threat to global health, with novel epidemic and pandemic viruses entering the human population from animal reservoirs [8–10]. Accurately identifying the host of a virus from its genome sequence alone is therefore a task with valuable practical and theoretical applications. Rapidly identifying the host of a novel virus has clear value in zoonotic disease surveillance and for the monitoring of high-risk animal reservoirs with sustained endemic transmission. Models can also be used to fill these knowledge gaps and infer the host origin of genomes from metagenomic samples. Computational prediction of viral traits therefore represents an important opportunity to address knowledge gaps that can provide actionable data to inform virological research and surveillance of potential zoonotic spillovers. In addition to these applications, the ability to predict viral host is dependent on underlying viral host co-evolution, and therefore enhanced understanding of the factors important for host prediction can shed light on underlying evolutionary mechanisms.

Virus host preference is a complex trait depending on numerous molecular interactions between a given virus and host. While some determinants of virus host range have been subject to detailed study in specific viruses, such as receptor binding [11,12], the function of many viral proteins and their interactions with host cell machinery remains unclear. Comprehensive mechanistic models of virus-host compatibility are therefore currently feasible only for a limited range of well-characterised interactions.

However, in the absence of detailed mechanistic frameworks, other information has been used to predict the hosts of specific viruses, including latent information in viral genomes. The coevolution of viruses with their hosts leaves distinct signatures in viral genomes [13,14]. Molecular evidence has revealed the mechanisms underlying some of these signatures. These include direct evolutionary pressure from host antiviral defences which induces adaptive changes in viral genomes to escape immune responses [15]. Viral use of host replication machinery also results in adaptation to optimise replication efficiency, for example by modulating codon usage in response to host tRNA availability [16,17]. Viruses may also mimic host short linear motifs (SLiMs), enabling interactions with host proteins which can be deregulated or hijacked to create favourable conditions for viral replication [18]. However, it is important to note that mimicry of host genome composition and conserved motifs is not the only adaptive pressure acting on genome composition [19]; the relationship between virus and host genome features is subject to a broader environment of constraints and cannot be defined in terms of straightforward mimicry. Machine learning models are well suited to learning these non-linear statistical associations between genome representations and host labels and thereby predict viral host directly from sequence data. A variety of machine learning models have seen broad application and success in making host predictions over a range of evolutionary scales and using different genomic feature representations [20–24].

However, these applications are subject to biases arising from a ‘triple collinearity’ between viral relatedness, host preference and genomic features. Because closely related viruses are likely to infect similar hosts and also have highly similar genome composition, predictive models can achieve strong predictive performance by functionally reconstructing viral phylogeny rather than finding true host-associated genomic motifs. Reliance on phylogenetic signal can limit the power of predictions for more divergent viruses and obscure which genomic features are genuinely associated with host adaptation. This makes elucidation of the mechanisms driving genome adaptation challenging. New modelling approaches that provide clear frameworks for reproducible and consistent benchmarking against phylogenetic prediction and interpretability of feature importance are therefore highly desirable, particularly as machine learning software packages are becoming more accessible to non-expert users.

We aim to provide such a framework through rigorous validation of host predictions at different phylogenetic scales. To disentangle the contribution of phylogeny from genomic features, we implement graph neural networks (GNNs) that explicitly encode viral phylogeny in the model architecture. We then benchmark model performance against predictions based on phylogeny alone across the same validation framework. Through this approach, we hope to clearly elucidate which elements of viral genomes contribute to host prediction through adaptation, rather than the proxy of phylogenetic signal.

GNNs are architectures that integrate relational structure with feature-based learning, enabling information from connected observations to inform prediction [25]. In biological applications, entities such as hosts, proteins or viruses can be represented as nodes connected according to known relationships. Information is then propagated between connected nodes through message passing, allowing each node’s representation to be informed by the context of nearby nodes in the graph. Graph representations are therefore ideally suited to the modelling of heterogeneously ordered data such as biological sequences. Decades of development of bioinformatic methods have provided powerful and accessible tools for determining the underlying phylogenetic structure of biological sequences. However, traditional supervised ML methods which are frequently applied to host prediction tasks typically make no use of this data. Explicit incorporation of viral phylogeny into the model architecture avoids requiring the model to learn phylogenetic relationships *de novo* every time a model is trained and may enable learning of adaptive signals with lower dimensional models and shorter training protocols. GNNs have been successfully applied to virus host prediction problems using a range of strategies[26–28], although this is the first application of GNNs which explicitly represent viral phylogeny as a graph to our knowledge.

We apply this methodology to assess the relative contributions of phylogenetic and adaptive signal to host prediction in the paramyxoviruses, a viral family of high priority for global disease surveillance. Paramyxoviruses are a family of negative-sense, single-stranded RNA viruses which includes numerous species of significance for human and animal health [29–34], including several emerging zoonotic threats that have been identified as high priority for pandemic surveillance, most prominently the henipaviruses [35,36]. A predictive tool that can assign viral host of origin with high confidence for paramyxoviruses can therefore enhance pandemic preparedness by identifying likely sources of spillover and high-risk human-animal interfaces.

In addition to their significance for global health, the paramyxoviruses make good candidates for the development of host prediction models due to their broad host range. Paramyxoviruses infect a wide range of vertebrates and these preferences are well-represented in sequence databases, allowing for relatively balanced host representation at different taxonomic ranks.

By applying GNN host prediction models to the *Paramyxoviridae* we elucidate the importance of phylogenetic representation to host prediction in a viral family which is a major target of zoonotic disease surveillance and metagenomic sampling [37–40]. Our thorough benchmarking and validation demonstrate the strong predictive power inherent to simple phylogenetic methods. Our novel GNN approach exceeds these benchmarks when making predictions on more divergent viral sequences, confirming that genuine signals of host adaptation are present in simple compositional representations of viral genomes, although they constitute a small proportion of overall predictive power when compared to the phylogenetic signal. We therefore present a model framework that can decompose these competing signals and assess importance of adaptive genome feature representations relevant for molecular host-virus interaction. This method can be applied to data on other host-virus systems and will enable more interpretable and accurately benchmarked approaches to host prediction.

## 2. Results

### 2.1 Data processing

We downloaded all available *Paramyxoviridae* genomes from NCBI Virus as of 21/11/2025 (n = 78,449). These sequences were cleaned and clustered to generate a set of centroid sequences representative of paramyxovirus diversity. Cluster-representative centroids were then used to extract the ORFs encoding 5 canonical paramyxovirus proteins: the nucleocapsid, matrix, fusion, attachment and polymerase proteins. BEAST X was used to generate a phylogeny of the representative sequences from these ORF sequences (Figure 1A). This phylogeny was used to construct a minimum spanning tree, with nodes representing individual viruses (cluster representatives) and edges scaled by patristic distance (Figure 1B). Compositional features were calculated from the extracted ORFs and at the whole genome level and assigned to the relevant node in the graph, in addition to host labels which were assigned at the taxonomic rank of order for mammals and class for non-mammalian vertebrates. The final graph contained 212 nodes representing 189 unique viral species, with 145 mammalian-infecting viruses, 60 avian-infecting viruses, four reptilian-infecting viruses and three fish-infecting viruses.

**Figure 1:**
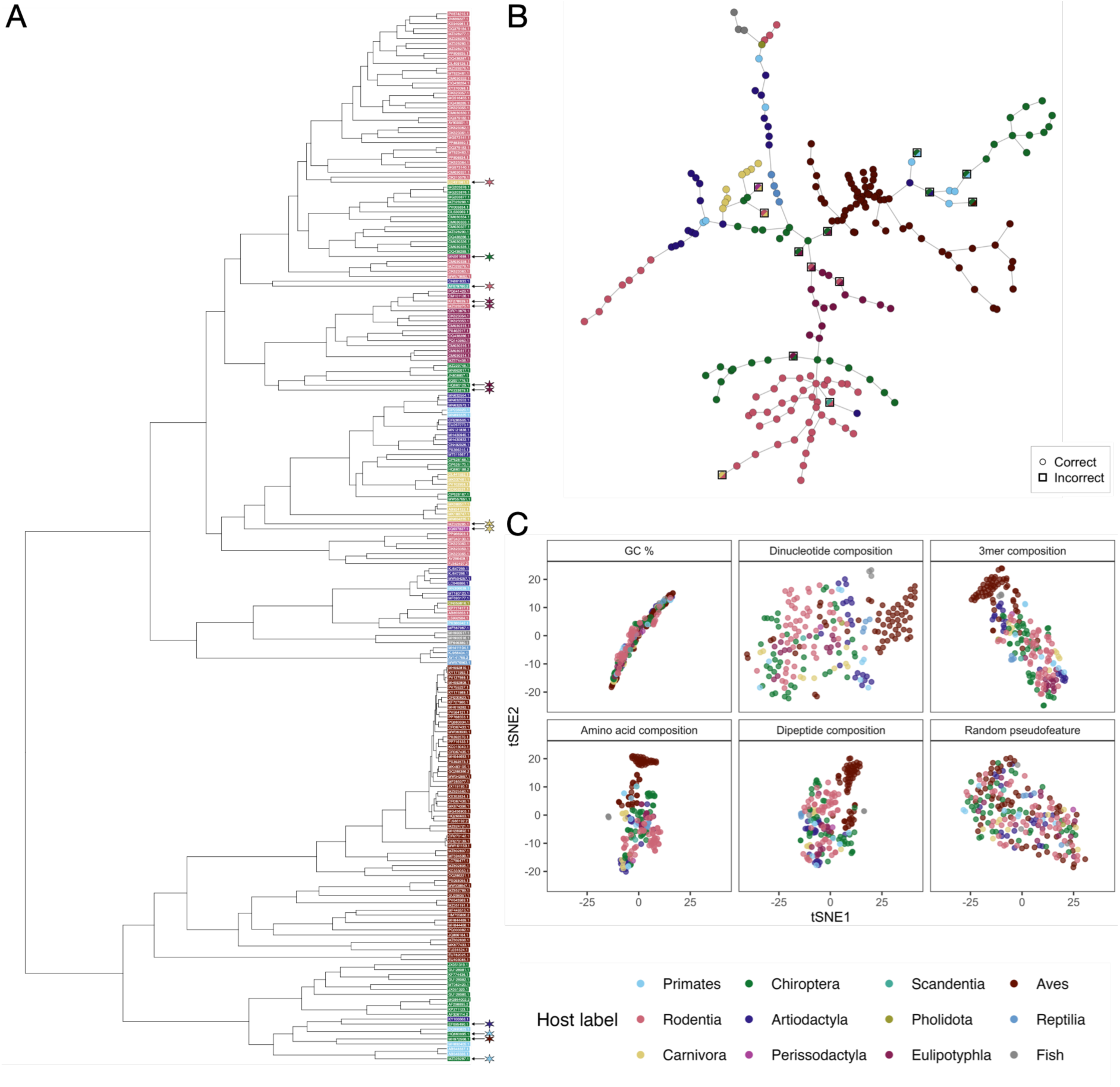
Figure illustrating degree of covariance between virus host, phylogeny and genomic composition and the results of host predictions made using BLAST. **A)** BEAST X phylogeny of representative paramyxovirus sequences. Tips are labelled with the sequence NCBI accession and coloured by host label. Asterisks mark sequences for which host was incorrectly predicted by BLAST, with the colour of the asterisk indicating the incorrectly predicted host label. **B)** Minimum spanning tree constructed from the phylogeny shown in A, with nodes coloured by host label. Nodes for which host was incorrectly predicted by BLAST are shown with black squares, with the coloured diagonal swatch in lower right indicating the incorrectly predicted host. **C)** tSNE embeddings of the five genome compositional features calculated and the random pseudofeature.

### 2.2 Assessing predictive power of phylogeny

We first aimed to understand the degree of covariance between host, viral phylogeny and viral genome composition (Figure 1A). Strong association between phylogenetic distance and host was evident (Mantel correlation statistic = 0.5322, p = 0.0001). t-distributed stochastic neighbour embedding (t-SNE) plots of the initial feature embeddings show clear clustering of virus host (Figure 1C), particularly for avian paramyxoviruses and the rodent jeilongviruses. Finally, a correlation of the Euclidean distances between raw feature embeddings and patristic distance shows that higher dimensional representations of genome composition are more representative of underlying phylogeny (Table 1), i.e. phylogenetic information pervades compositional feature representations.

**Table 1:** Table calculated compositional features, their dimensionality and their distance from the phylogeny. Note that dimensionality values are given for full feature sets calculated from five ORFs and the whole genome (nucleotide features).

| Compositional feature | Dimensionality | Sum squared residuals |
| --- | --- | --- |
| GC % | 6 | 20346 |
| Amino acids | 100 | 9239 |
| Dinucleotides | 288 | 11661 |
| 3mers | 384 | 8713 |
| Dipeptides | 2000 | 5470 |

In order to quantify the predictive contribution of phylogeny and genome adaptation, we first benchmarked the host-predictive potential of phylogeny alone. Our primary phylogenetic benchmark used the BLAST algorithm, with separate queries made for each of five extracted ORFs and a final prediction made by majority vote across the five queries. We also conducted additional filtering of highly skewed results, resulting in 92% accuracy across the full dataset, compared with 84% without additional filtering of skewed hits. Incorrect predictions were distributed across the phylogeny as shown on both the tree and graph representations (Figures 1A, 1B), with incorrect predictions frequently made on viruses with long branch lengths or in viral subclades which infect a range of hosts.

### 2.3 Model initialisation

We next compared GNN performance with similarly sized shallow feed-forward neural networks (FNNs) and random forests (RF) across the different compositional feature sets. Both GNNs and FNNs reached a maximum accuracy of 98% using the dipeptide compositional feature (Figure 2A-B), while the BLAST benchmark and RF achieved 92% accuracy.

**Figure 2:**
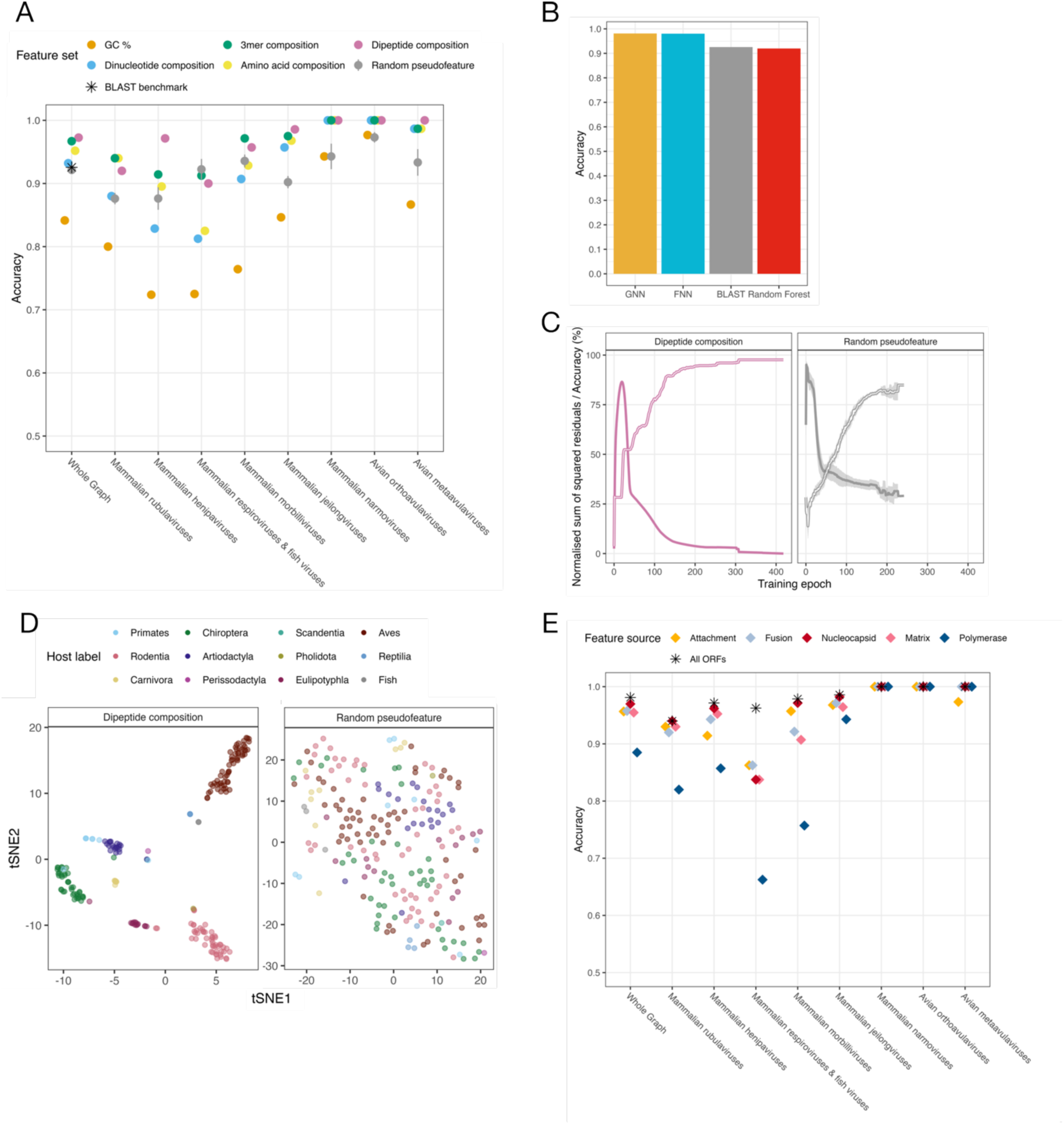
**A)** Performance of different feature sets across the whole dataset (leftmost) and taxonomically relevant subgraphs. The asterisk shows the performance of the BLAST benchmark. Error bars show ±1 standard deviation across the five randomly initialised pseudofeatures. Subgraphs with fewer than 5 cluster representatives are excluded from the plot. **B)** Maximum achieved accuracy using four predictive methods: GNN, FNN, phylogenetic inference with BLAST and random forest. **C)** Model accuracy (hollow line) and normalised mean sum of the squares of residuals (solid line) between patristic distance and Euclidean distance in embedding space over model training epochs, for models using dipeptide compositional features and the random pseudofeature. The solid line shows the residual, and the hollow line shows accuracy. Ribbon shows ±1 standard deviation across the five randomly initialised pseudofeatures. **D)** t-distributed stochastic neighbour embedding projection of embeddings following GNN model training with dipeptide compositional feature or random pseudofeature. **E)** GNN model performance using dipeptide compositional feature calculated using 5 different ORFs across the whole dataset (leftmost) and taxonomically relevant subgraphs. Asterisks show the performance of models trained using concatenated features from all 5 ORFs.

To test how much predictive performance could be recovered from graph structure alone, we trained models using random pseudofeatures. These models achieved ∼92% accuracy, closely matching the BLAST benchmark (Figure 2A), whereas the corresponding FNN reached 79%. We next defined a set of taxonomically relevant subgraphs (Table 2) to identify whether predictive power varied across subclades. Analysis of performance over subgraphs showed very strong ability to distinguish paramyxoviruses infecting avian hosts, with 100% accuracy for most feature sets including GC % despite this feature being largely uninformative for predictions of other host clades. The weakest performance was on a highly diverse clade representing the genus *Respirovirus*, which infect a range of mammalian orders, as well as all paramyxoviruses which infect fish. No feature set outperformed the random pseudofeature in this subgraph.

**Table 2:** Table showing taxonomic representation, primary host and number of members of the defined subgraphs.

| Viral clade represented | Primary host clade | Number of members |
| --- | --- | --- |
| (whole graph) |  | 212 |
| *Rubulavirus* | Mammals | 20 |
| *Henipavirus* | Mammals | 21 |
| *Respirovirus* and related | Mammals and fish | 16 |
| *Morbillivirus* | Mammals | 28 |
| *Jeilongvirus* | Mammals | 56 |
| *Narmovirus* | Mammals | 7 |
| *Orthoavulavirus* | Birds | 43 |
| *Metaavulavirus* | Birds | 15 |
| *Ferlavirus* and related | Reptiles | 4 |
| *Paraavulavirus wisconsinense* | Birds | 2 |

Under optimal tuning parameters, all compositional features (excluding GC%) resulted in strong predictive performance. However, higher dimensional features were generally more robust to different tuning conditions, with dipeptide composition maintaining high accuracy across a range of hyperparameter conditions (Supplementary Figure 1, Supplementary Figure 2). We therefore selectively show results from models trained using the dipeptide compositional feature hereon. We also constructed an alternative phylogeny using only the ORF coding for the RdRp. Using this phylogeny resulted in lower overall predictive performance, confirming that our model pipeline was suitable for the prediction task (Supplementary Figure 3).

To further interrogate how much model-learned embeddings reflected phylogeny, we directly compared two pairwise distance matrices: patristic distance, and Euclidean distance between learned model embeddings. We expect that as the model learns, embeddings will be updated to more closely represent underlying phylogenetic distances and residuals between both distance matrices would reduce. Figure 2C supports this hypothesis. Residuals decreased during training for both feature sets, although dipeptide-based models showed an initial increase associated with separation of avian and non-avian viruses before declining after epoch 19. Final residuals were substantially lower for dipeptide features than for the random pseudofeature (unnormalized sum square residual values of 5033 vs 10110), indicating a closer correspondence with phylogenetic structure. The final learned model embeddings are shown projected in 2 dimensions with t-SNE in Figure 2D, clearly showing the weaker clustering of host labels for models trained using the random pseudofeature relative to dipeptide composition.

We finally explored how calculating features from different genes affected model performance. Figure 2E shows performance over the subgraphs defined in Table 2 with compositional features calculated from each of 5 ORFs. Features calculated from the L ORF, encoding the RdRp, result in weaker predictive performance across most subclades, although these features still distinguished avian vs non-avian infecting viruses, and the difference was less pronounced for the jeilongviruses. Features derived from the ORF encoding the nucleocapsid protein meanwhile slightly outperform features from other ORFs in most subclades. For most subclades the accuracy achieved training models using features derived from nucleocapsid alone approach the accuracy achieved using features from all 5 ORFs; however, for the respiroviruses a combination of features from multiple ORFs was essential to achieve good prediction accuracy.

### 2.4 Exploring model performance over different scales

We next assessed model generalisability across increasing evolutionary distances using two extrapolative validation strategies relevant to real-world surveillance (Figure 3). Leave-one-out (LOO) validation tested short-range extrapolation by withholding individual nodes, whereas blocked cross-validation tested prediction of more divergent viruses by withholding entire phylogenetic subclades defined in Table 2.

**Figure 3:**
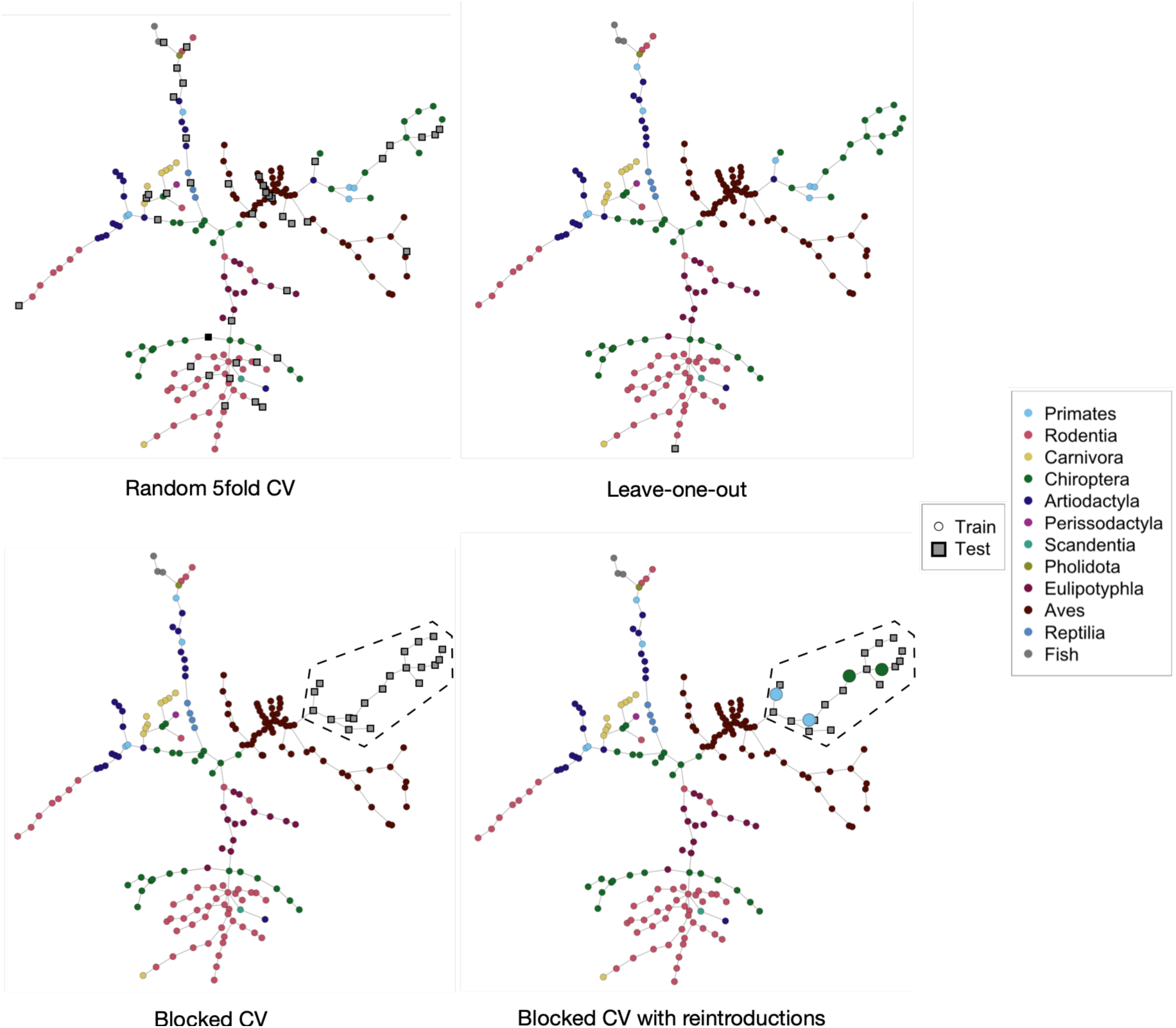
Representations of different validation strategies. Nodes in the training set are represented as circles coloured by host label. Nodes in the test set are represented as grey squares with black outline. Top left graph shows representative random 5-fold cross-validation. Top right graph shows representative LOO validation with a single node held out. Bottom left graph shows representative blocked validation, with the rubulavirus subgraph shown blocked (highlighted with dashed line). Bottom right graph shows representative blocked validation with 4 random nodes reintroduced into the training set from within the rubulavirus subgraph (shown double scale).

In order to maintain a consistent and comparable benchmark, we also generated BLAST query databases in which equivalent held out data was removed, albeit BLAST uses all sequences in the dataset and is not limited to cluster centroids.

LOO validation resulted in similar performance between predictions made with both NN models and phylogenetic predictions made with BLAST. GNN and FNN predictions made using the dipeptide composition feature correctly predicted 190 of 212 nodes (89.6%), while BLAST predictions correctly assigned 189 (89.2%). Of these incorrect predictions, the majority (n = 19) were shared between the two methods (Figure 4A). In addition, those nodes for which BLAST was incorrect and the GNN was correct tended to be lower confidence predictions, suggesting that similar sequences are challenging prediction targets for both methods.

**Figure 4:**
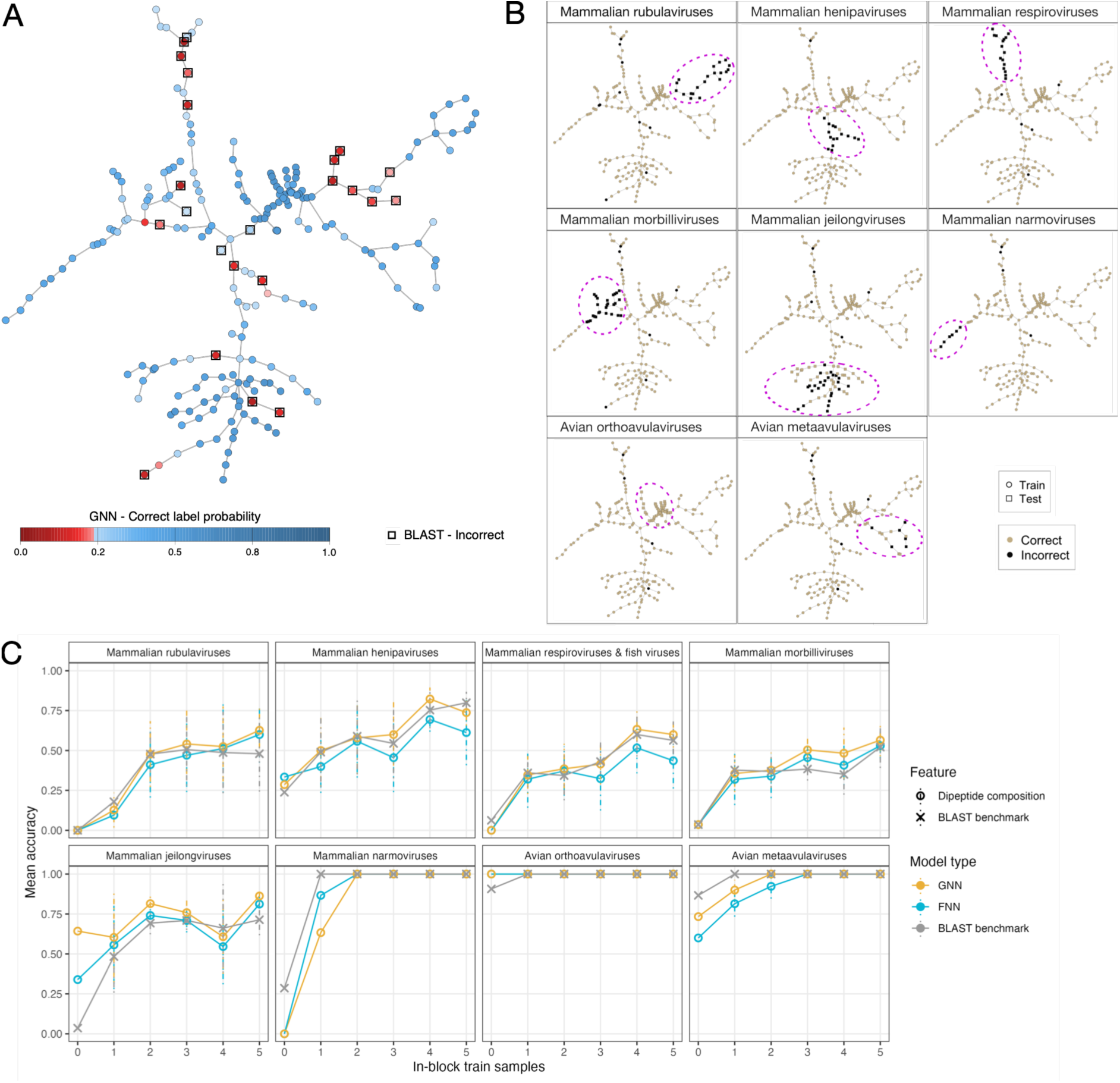
Figure showing model performance with LOO and blocked validation strategies. **A)** Predictions made by the GNN dipeptide composition model with a LOO validation strategy. Blue nodes indicate correct predictions and red nodes indicate incorrect predictions, with colour intensity showing the confidence of each prediction. Nodes incorrectly predicted by the BLAST benchmark are indicated with black squares. **B)** Prediction across blocked cross-validation folds. The blocked subgraphs are indicated by node shape (circular nodes indicate train nodes, square nodes indicate test nodes) and are additionally highlighted with a magenta dashed line. **C)** Blocked cross validation performance of GNN and FNN models and the BLAST benchmark following reintroduction of nodes from within the blocked subgraphs. Dashed error bars show ±1 standard deviation across 5 randomly selected reintroductions.

However, the more stringent blocked cross-validation approach showed an advantage for the GNN model over both BLAST predictions and predictions made with the FNN. Most of this increased performance was driven by the *Jeilongvirus* clade. When *Jeilongvirus* sequences were entirely held out of the reference database, BLAST uniformly predicted *Eulipotyphla* hosts for all nodes in this subgraph, resulting in 4% accuracy. In contrast, the GNN correctly distinguished many *Jeilongvirus* sequences from rodents and bats, achieving 64% accuracy (Figure 4B), while the FNN achieved an intermediate accuracy of 34%. We additionally tested reintroducing sequences from within the blocked subgraph to identify how many more sequences from within a subclade are needed to recover predictive performance. Nodes from within the blocked subgraph were selected at random and between one and five nodes were included in the training set (Figure 3). The random selection was repeated five times. Predictive performance was greatly improved for all predictive methods in some subgraphs by the presence of only one or two sequences from within the blocked clade in the training set (Figure 4C). GNN models showed a small but consistent improvement in accuracy over FNN models for all the mammalian-infecting paramyxovirus clades following reintroduction of sequences.

As a final performance test, we predicted the host of paramyxovirus sequences added to NCBI Virus after initial data extraction (21/11/2025). These were subject to an identical cleaning and clustering pipeline before being added to the existing tree based on their RdRp sequence using a maximum likelihood approach in IQ-TREE 3. The new sequences represented 39 unique viral species (of which 6 were not present in the original training set) which were well-distributed over the phylogeny (Figure 5), providing a good case study in application of the model to real-world surveillance tasks. GNN models trained on the original graph were then used to make predictions on these new sequences and BLAST was again used to generate a phylogenetic benchmark. BLAST correctly predicted the host of 51 out of 56 (91.1%) new sequences, while the GNN and FNN models exceeded this, correctly predicting the host of 55 (98.2%; Figure 5). The only sequence incorrectly classified by the NNs was a highly divergent bearded seal-associated virus [41]. This has very limited sequence conservation with any known virus sequence and is only tentatively classified in family *Paramyxoviridae*. The true label of this sequence is in doubt; while isolated in a bearded seal (and therefore was given ground truth host label *Carnivora* for our analysis) it may have entirely different host(s) as a possible dietary contaminant [41].

**Figure 5:**
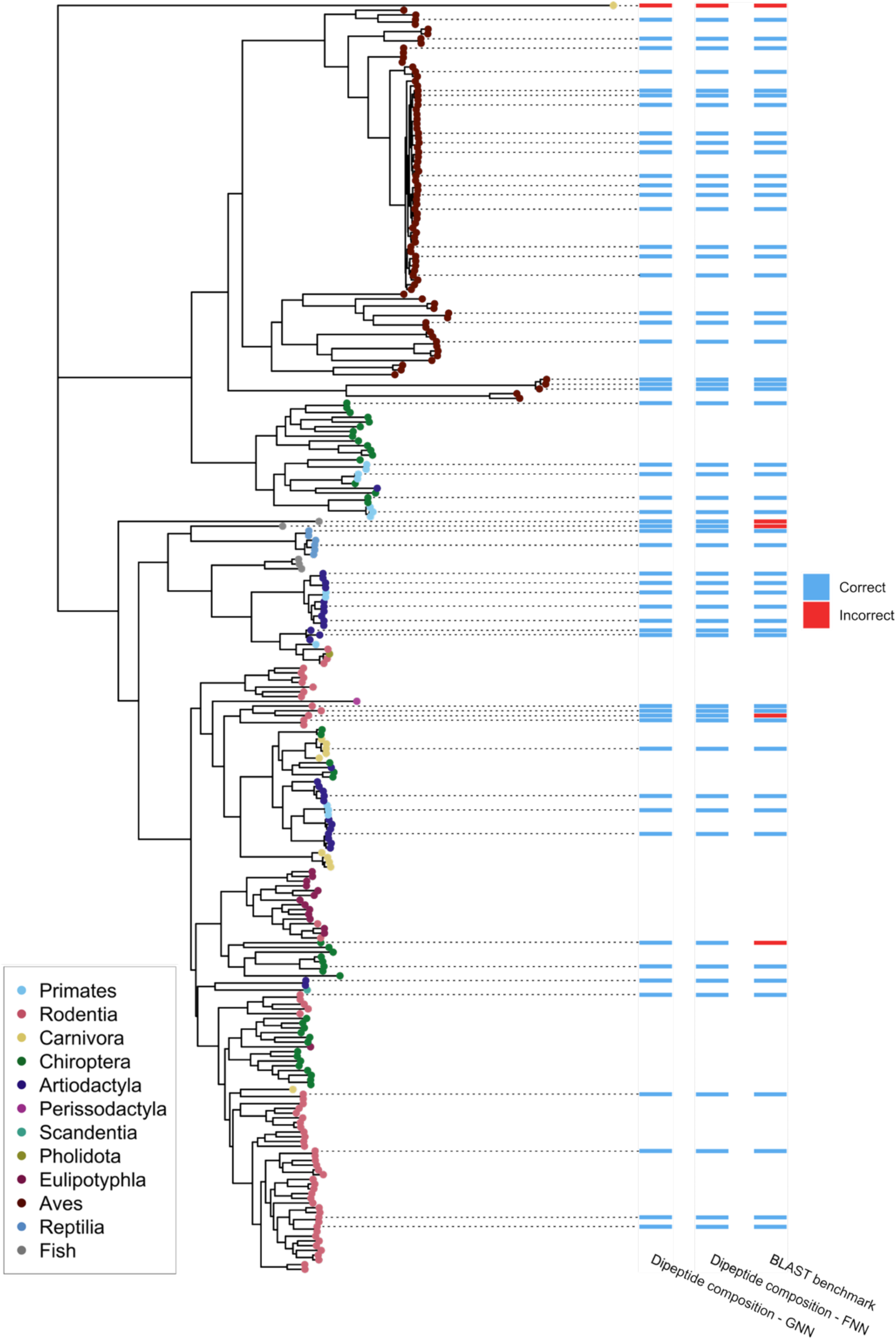
Phylogeny and predictive performance on 55 recently published out of sample paramyxovirus sequences. The phylogeny representing the relationship between all sequences in the training set and the new out of sample sequences is shown on the left-hand side. Heatmap indicates the predictions made by different prediction methods for the new sequences.

Overall, phylogenetic inference can generally make sufficiently accurate predictions of the host of origin of a virus when closely related viruses are represented in the reference database. However, predictive power declines for more evolutionarily divergent viruses, where models that incorporate genomic features provide a clear advantage. Neural networks outperform phylogenetic inference across longer evolutionary distances, with GNNs showing the strongest performance when predicting entirely unseen subclades.

## 3. Discussion

### 3.1 Performance of supervised learning models

GNNs demonstrate advantages for the prediction of virus-host associations. The explicit representation of phylogeny in the model provides additional context which is highly relevant to the host prediction task. In non-phylogenetically structured models this context must be learned *de novo* from genome feature representations. This results in clear advantages for GNNs in predictive tasks where no closely related sequences are present in the training data. The superior performance of GNNs during blocked CV (Figure 2C) suggests that message passing between related nodes helps retain host-relevant information when training data is sparse. The particularly strong performance of the GNN on the *Jeilongvirus* subclade is more difficult to explain. Jeilongviruses encode additional proteins and possess several genomic features distinct from other paramyxoviruses [42,43]. Notably, L-ORF-derived features performed similarly to those from other ORFs in this clade, unlike in most other subclades (Figure 2E). This suggests that jeilongviruses may possess distinctive genome-wide compositional signals that are captured effectively when combined with phylogenetic context. In the blocked-validation setting, message passing may then allow these embeddings to be shifted towards a region of feature space associated with the dominant *Jeilongvirus* host orders, *Chiroptera* and *Rodentia*.

However, in many cases FNNs or even simple phylogenetic inference using BLAST provide equally strong predictive performance. For host prediction from viral sequences closely related to viruses already represented in large public databases, a simple BLASTn search performs as well as our best models, as demonstrated by LOO validation. Where NN models do show an advantage over purely phylogenetic approaches such as BLAST is in more extrapolative cases; making predictions on new virus clades as demonstrated by the blocked cross-validation results or over long branch lengths as demonstrated by the performance on two unseen sequences from divergent fish-infecting viruses Hippocampus erectus paramyxovirus 1 and *Aequorvirus hippoglossi* (Figure 5). For metagenomic applications or predictions in viral families with little existing representation in sequence databases GNNs are likely to provide superior predictions to other approaches.

### 3.2 Feature sets

We applied these models using simple compositional features of genomes. We deliberately focused on simple compositional features because the number of non-redundant viral sequences with confidently assigned hosts was insufficient to support more complex feature representations. While this work clearly shows that simpler compositional features do encode some host information in addition to phylogeny, it is likely that we are approaching the limits of how much information can be extracted from such simple representations of viral genomes. Recent advances in machine learning such as the application of large language models to biological sequences promise feature representations better able to capture the high dimensional underlying biological traits that underpin host preference [44,45]. Future applications making use of both phylogenetically structured model architectures and language model-based features are an excellent prospect for further enhancing the performance of host prediction models, although space may remain for simpler feature representations which foster greater interpretability and inference of underlying adaptive evolutionary processes.

### 3.3 Benchmarking and validation of host prediction models

Model development was conducted using 5-fold cross-validation. This validation strategy is unlikely to give a complete picture of performance when working with highly structured data, including phylogenetically structured data [46]. Due to random partitioning of graph nodes into train and test folds, nodes in the test set are typically proximal to a node featured in the train set. In our dataset, 91.5% of test nodes have an immediate neighbour which is in the training set and 7% are two edges from a training node, while no test nodes were greater than 4 steps from a node in the training set across the 5 CV folds used to assess model performance. K-fold cross validation would therefore give a good estimate of interpolative performance (performance within the existing range of the data) but not necessarily a good estimate of extrapolative performance. Genome sequencing has captured only a small proportion of vertebrate viral diversity and viral datasets are therefore very sparse; it is therefore essential that predictive models intended for use in surveillance or metagenomic sampling scenarios apply non-random blocking for validation if they are to be generalisable. Our approach depended on graph connectivity metrics, with thresholds selected to give phylogenetically relevant subclades. Other approaches include clustering by distance in a feature space [47]; this approach could be considered even more rigorous as blocking can be carried out using distances derived from the same features as model inputs. Our results clearly demonstrate the importance of thorough validation with a range of strategies, as the benefit of GNN over FNN architectures were only evident in blocked validation applications (Figure 4).

In addition to validation, this work demonstrates the importance of thorough phylogenetic benchmarking. Achieving apparently strong predictive performance for host prediction tasks using supervised learning is trivial given the strong collinearity between phylogenetic signal and viral host of origin. It is essential that new trait prediction models are benchmarked against a phylogenetic predictor. Neglecting comparison to phylogenetic inference undermines the claims of performance made about specific feature sets and model architectures, and risks leading to the prioritisation in future research of sub-optimal methodologies. Previous work has attempted to identify the contribution of phylogeny to host prediction in various ways; this includes phylogenetic benchmarks using BLAST similar to the benchmark applied in this paper and comparisons of dendrograms derived from phylogeny to those derived from hierarchical clustering of feature importance [20,45]. It is notable that we saw a large improvement in the accuracy of the BLAST benchmark through a simple process of running searches across multiple ORFs and filtering based on the skewness of the top hits (92% with skew correction vs. 84% without skew correction). Taking steps to ensure the optimal hits returned from a BLAST search are used for host determination can lead to substantial improvements in results but are frequently neglected, in turn leading to overestimation of the adaptive signal present in genome feature representations.

### 3.4 Paramyxoviridae dataset

A key motivation to conducting this modelling on a limited dataset was to ensure accuracy of manual host label assignment. We aimed to describe features which optimally represent the result of co-evolution between virus and their hosts. Many public databases of virus-host range assign host based on sample source from sequence metadata which may represent cross-species spillovers rather than long-term evolutionary relationships. The uncertainty around host status has led such data to be conservatively termed host-virus association [27]. Predictions made on these datasets are also addressing an important, and more challenging, question: that of host permissibility, the detection of which will depend on different signals to prediction of the host of origin. Host-prediction datasets should therefore distinguish long-term evolutionary associations from transient infection or exposure, as these relationships are likely to be encoded by different biological signals. Indiscriminate use of data where transient infections and long-term co-evolutionary relationships are conflated will result in the obfuscation of signals specific to these different types of relationships.

Our results show that explicitly incorporating phylogeny improves host prediction when models are required to extrapolate beyond closely related reference sequences. More importantly, they show that high predictive accuracy alone is insufficient evidence of host-associated genomic adaptation unless performance is benchmarked against phylogeny. Future host-prediction models should therefore combine phylogenetically informed architectures, rigorous extrapolative validation and explicit phylogenetic baselines if they are to identify genuinely informative genomic signals.

## Methods

### Nucleotide processing

All whole genome sequences in the family Paramyxoviridae uploaded to NCBI virus prior to 21/11/2025 were downloaded. The dataset was filtered to remove sequences with >1% ambiguous positions and those within the first quintile of total sequence length. Several other transgenic or low-quality sequences were removed; a full list of filtered sequences is available in the dropped_sequences.csv file in Supplementary Information. All sequence processing was carried out in R with the biostrings package [48] or using the seqkit2 command line utility [49].

This set of clean nucleotide sequences was then clustered using the Mmseqs2 Linclust algorithm [50] to reduce over-representation of densely sampled viral species while preserving sequence diversity across the family. A grid search was performed over the minimum sequence ID and minimum coverage hyperparameters. Optimal values for these parameters were determined by inspecting several metrics, comprising total number of clusters, number of viral species represented in each cluster, number of replicate centroid sequences per viral species and a silhouette score used as assessment of cluster quality or ‘tightness’. For the cluster quality assessment, the pairwise Jensen-Shannon distance between the 3mer profiles of every pair of sequences was calculated; these pairwise distances were used to compute the silhouette score. Consideration of all of these metrics resulted in selection of final hyperparameters for the clustering algorithm of minimum sequence ID = 0.9 and minimum coverage = 0.4. This gave 218 clusters in total covering 191 unique viral species. The cluster centroid sequences were used for all downstream applications.

### Host assignment

Following clustering, we assigned a host label to every cluster. Host label was selected by manual review to reflect the host origin; that is, the long-term host within which we expect virus sequences in this cluster to have co-evolved. Sequences are therefore anticipated to contain adaptive signatures resulting from the evolutionary relationship with their host labels.

We established a virus-host lookup table for every virus species in the dataset. To generate this lookup table, we first converted the names of host species listed in sequence metadata into taxonomic IDs using the taxizedb package in R [51], and thereby assigned hosts at the taxonomic class and order ranks. For cases where host metadata was absent (n = 4), we conducted a literature search and manually assigned those clearly identifiable. In cases where there was unanimous agreement between all sequence representatives of a viral species on its host taxon, that taxon was assigned in the lookup table. In cases where there was not unanimous agreement between all sequences on the host taxa, we conducted a literature search and manually assigned a consensus taxon to the lookup table (Supplementary Table 1).

A final host label for each cluster was then determined based on this lookup table. For mammalian-infecting viruses, the host label was at the order rank (either Primates, Rodentia, Chiroptera, Artiodactyla, Perissodactyla, Carnivora, Eulipotyphla, Pholidota or Scandentia). For non-mammalian infecting viruses, the host label was at the class rank (either Aves, Reptilia, or a pseudo-taxon Fish which grouped five viruses infecting organisms in the subclass Actinopteri and one virus infecting the class Chondrichthyes). Two species of mammalian-infecting viruses could also not be assigned at the level of taxonomic order; mammalian orthorubulavirus 5, a generalist that readily infects mammals from many orders, and a novel morbillivirus from Brazil whose cluster contained a centroid sequence sampled from bats but several peripheral sequences sampled from marmosets. Clusters representing these viral species were dropped, resulting in 215 final centroid sequences.

### ORF extraction and validation

Detection of open reading frames was carried out using the R package ORFiK. We aimed to designate 5 canonical ORFs for all paramyxovirus genomes to ensure consistent features could be calculated. These were N, M, F, H/HN/G and L, which encode the nucleocapsid, matrix, fusion, attachment and polymerase proteins respectively.

Cluster centroid genomes were first split by whether they belonged to a jeilong-like virus, which have longer genomes with distinct architecture compared to most paramyxoviruses. A rules-based ORF detection regimen was then employed, which designated ORFs based on relative length and position in the genome. Different rules were applied to jeilong and non-jeilong viruses. Where ORFs could not be assigned using the rules-based detection, or replicates of the same ORF were detected, the correct designation was determined by pairwise alignment to a reference sequence. Where a reference sequence for the same viral species was not available, the reference sequence of a related virus was used. Finally, the detected ORFs of all cluster centroids for which a reference sequence belonging to the same viral species was available were aligned to the reference to validate the ORF detection process. Any alignments with a negative substitution score (as determined by the pairwiseAlignment function in the pwalign R package [52]) were manually inspected to ensure the detected ORFs had been correctly assigned. This allowed for 132 of 215 total centroid genomes to be validated against a RefSeq. All validated sequences were clearly homologues of the reference, confirming that the ORF detection process was reliable.

### Feature generation

Calculated nucleotide features were GC %, dinucleotide composition and 3-mer composition. Nucleotide features were generated for the whole genome and the five identified canonical ORFs for each cluster centroid sequence. 3-mer counts were calculated using overlapping windows with a one-nucleotide step.

Calculated protein features were amino acid composition and dipeptide composition. Protein features were generated for the five identified canonical ORFs for each cluster centroid sequence. Prior to generation of amino acid features, identified ORFs were translated into amino acid sequences. We attempted to repair ambiguous bases which resulted in ambiguous amino acids in the protein sequence by aligning the cluster centroid to other sequences belonging to the same cluster. A consensus matrix was then calculated at the position of the ambiguous nucleotide and the most frequently occurring nucleotide taken as the consensus. Where a consensus could not be generated due to lack of additional sequences from the same cluster or lack of consensus across sequences, the amino acid was dropped.

All compositional features were calculated as frequencies and normalised by sequence length.

Random pseudofeatures were generated with the same dimensionality as the smallest feature set GC %, i.e., a six-dimensional vector (one GC % value for each ORF in addition to one GC % value for the whole genome). The random numbers were sampled in the same min-max range as real GC % values but were not constrained to the same distribution, resulting in a uniform distribution. Five random pseudofeatures were generated and used for all validation strategies and modelling approaches. Separate models were trained on each of the five random pseudofeature sets with final performance calculated as the mean accuracy across these five models with error bars given as ±1 standard deviation.

### Phylogeny

To generate a phylogeny, a multiple sequence alignment (MSA) was created for each of the five identified canonical ORFs from all centroid sequences using MAFFT [53] in global (G-INS-i) alignment mode with two maximum iterations.

ModelFinder [54] was used to identify the best-fitting nucleotide substitution model for each alignment according to Bayesian information criterion. The general time-reversible (GTR) model was selected as the optimal substitution model for all individual ORF partitions.

The five ORF alignments were then analysed jointly in BEAST X [55], with a shared tree topology and substitution-model parameters estimated independently for each partition. Rate heterogeneity among sites was modelled using a gamma distribution with dour discreet rate categories for each ORF. A constant-size coalescent model was used as the tree prior.

Alternative molecular-clock models were evaluated in preliminary analyses, including Hamiltonian Monte Carlo (HMC) relaxed clock, uncorrelated lognormal relaxed clock and a strict clock. Preliminary clock model analyses were run for 10 million MCMC states. Model fit was compared using path-sampling/stepping-stone marginal likelihoods, and the HMC relaxed-clock model was selected for final analysis (Supplementary Table 2).

The final analysis used partition-specific GTR+Γ4 substitution models and HMC relaxed clocks and was run for 50 million MCMC states, with parameters and trees sampled every 10,000 states. The posterior tree distribution was summarised using HIPSTR to obtain the final phylogeny used for downstream analysis.

To generate a graph from the phylogeny, patristic distances were calculated between all sequences in the tree. The graph was constructed with individual sequences as nodes connected by edges weighted by the patristic distance. The minimum spanning tree was then calculated based on these edge weights and used for model training.

Subgraphs were defined by thresholding the weighted minimum spanning tree with a patristic-distance cut-off and treating the resulting connected components as separate subgraphs [56]. The cutoff was selected to yield phylogenetically coherent subgraphs corresponding to taxonomic groups shown in Table 2, with a final cut-off selected as the 33^rd^ centile of the mean edge weight (edges ≤ 37.2 were dropped).

### Neural network model training

Graph neural network models were built in Julia using the GraphNeuralNetworks.jl package [57]. Graph models consisted of a different number of hidden GCN layers [58] using Rectified Linear Unit (ReLU) activation functions, with a final fully-connected classification layer of dimension size 12 representing the 12 host label classes. Where models were trained with dropout, this was applied before the final layer. Cross-entropy was used as model loss and optimisation was performed with Adaptive Moment Estimation (Adam). The update of node *i* is updated by aggregating messages from neighbouring nodes *j* ∈ *N*(*i*):

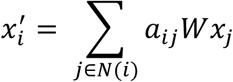

where *W* is a trainable weight matrix and *α_ij_* is a normalisation function used to weight messages by edge weights:

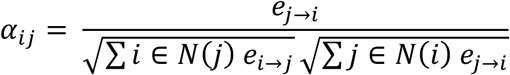

where the edge weights *e_i→j_* and *e_j→i_* are given by the reciprocal of the patristic distance between node *i* and *j*. The model was trained without self-loops and edges were treated as undirected, with weights defined as the reciprocal of patristic distance between connected nodes.

Five-fold cross-validation folds used to train initial models were generated using the createDataPartition function in the R package caret, which assigns observations to folds so as to optimally balance classes across folds. Hyperparameter tuning was conducted for models using each of the 10 feature sets (5 compositional features + 5 random pseudofeatures). A grid search was conducted over models using 2, 3 or 4 hidden convolutional layers with a dimension size of the hidden layers of 32, 64, 128, 256, 512 or 1024. The combination of tuning parameters that resulted in optimal accuracy for each feature set was determined and for all subsequent tests these optimal tuning parameters were used. The modal optimal tuning parameters were selected across the five different random pseudofeatures and used for all subsequent models using the random pseudofeatures. Model evaluation as presented in Figure 2B was performed on a different set of random folds to those used in model selection and parameter tuning.

An early stopping function was used to end training if model loss had reached an equilibrium or was increasing. At each training epoch the slope *β* was estimated by linear regression of model loss against epoch number over the preceding 10 epochs. If 0 ≤ *β* then model training was ended. For larger models (number of hidden layers ≥ 3 and dimension size of hidden layers ≥ 512) *β* was instead calculated over the preceding 30 epochs and additionally it was required that model loss had increased for each of the preceding 10 epochs before training was ended. This prevented short term fluctuations in loss resulting in premature stopping of model training.

To provide a non-graph structured comparison to the GNN models, we trained shallow feed-forward models using the flux.jl package [59]. FNN models consisted of a different number of hidden layers using ReLU activation functions, with both number of layers and dimension size of the hidden layers tuned with an identical approach to that employed for GNNs. Dropout was likewise employed before the final classification layer, and FNN models were trained with the same Adam optimiser, loss function and early stopping criteria as the GNN models.

#### Blocked cross validation and reintroduction

Non-random validation was carried out by holding out every node within each of the defined subgraphs and training a model on the remaining graph. Predictions were then made on every node within the blocked subgraph and average accuracy calculated across these predictions. For blocking with reintroduction, between one and five nodes from within the subgraph were selected at random and included in the training set. Predictions were then made on the remaining subgraph nodes not included in the training set. This random sampling was repeated five times and performance calculated as mean accuracy, with error bars given as ±1 standard deviation.

#### Out of sample sequences

All sequences used for model development and validation were deposited prior to 21/11/2025. To generate an out of sample test set we downloaded all Paramyxoviridae sequences deposited after this cutoff date and processed them using the same pipeline. In order to incorporate them into the existing phylogeny, a multiple sequence alignment of the identified L protein (RdRp) ORFs of both the old and new cluster centroids was computed using MAFFT as described above. IQ-TREE 3 [60] was then used to add the sequences based on this alignment, using the existing BEAST X tree as a constraint tree. This replicates a real-world surveillance application in which rapid prediction of putative host of a novel virus is required.

### BLAST benchmarks

BLAST benchmarks were conducted by generating a custom database containing all the sequences belonging to any cluster for which the cluster centroid representative sequence was in the training set for a given validation strategy. The nucleotide sequences of the five extracted canonical ORFs were then queried against this database using Blast+ with the discontiguous megablast algorithm. Multi-stage filtering for closest matches was conducted. Initially hits in the top 1% bitscore and bottom 1% E value within the given query were extracted. We then calculated the skew of the bitscores of the extracted hits and performed additional filtering depending on the severity of the skew, as shown in Table 3. This ensured only the closest sequence matches were used in BLAST host predictions. The modal host prediction was calculated for each ORF query and a final prediction made by majority voting across the five ORFs. In cases where votes were tied, the host with the highest summed bitscore across final sequence hits was selected.

**Table 3:**
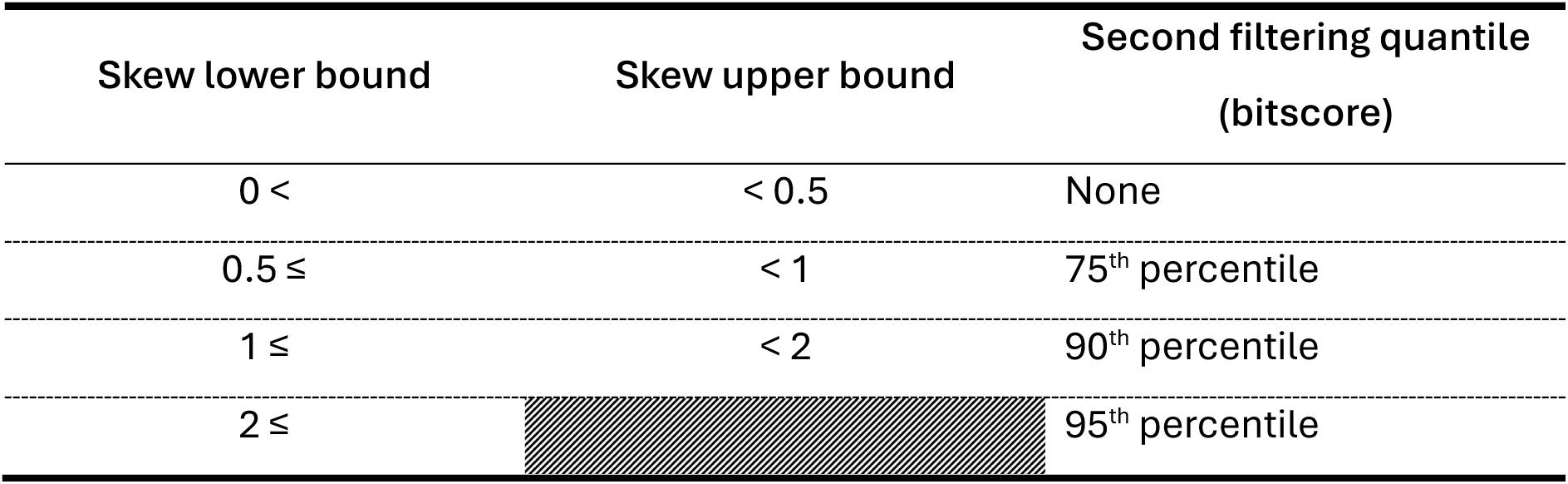
Skew limits used to conduct second round of match quality filtering for BLASTn host prediction.

| Skew lower bound | Skew upper bound | Second filtering quantile<br>(bitscore) |
| --- | --- | --- |
| $0 <$ | $< 0.5$ | None |
| $0.5 \leq$ | $< 1$ | 75<sup>th</sup> percentile |
| $1 \leq$ | $< 2$ | 90<sup>th</sup> percentile |
| $2 \leq$ | | 95<sup>th</sup> percentile |

### Random forest benchmarks

Random forests were trained using the ranger package in R [61]. Hyperparameter tuning was carried out by grid search across minimum node size (tree depth) of 1, 5 or 10 and total number of features considered at each split of 5%, 10%, 20% or 50% of the total number of features in each set. Splitting rules were set to gini and extratrees. Performance was calculated as mean accuracy across the same five cross-validation folds used to assess performance of neural net models.

### Model embedding-phylogeny residual

To assess the degree to which model embeddings were updated to reflect underlying phylogenetic relationships, we compared embedding and phylogenetic distance matrices. At each epoch, pairwise Euclidean distances were calculated from the final hidden-layer node embeddings. The sum of squared differences between the embedding-distance matrix and patristic-distance matrix was then calculated across all unique node pairs. Residual values were min-max normalised to 0-100 separately within each training run for visualisation.

## Data availability

All data and scripts are available on GitHub at: anonymous.4open.science/r/GNN-host-prediction-5131

## Acknowledgements

The authors thank Joseph Hughes for valuable early feedback on this project, Paul Johnson for advice on validation strategies and Jordan Bone for support with BEAST X.

## Supplementary Tables

**Supplementary Table 1:**
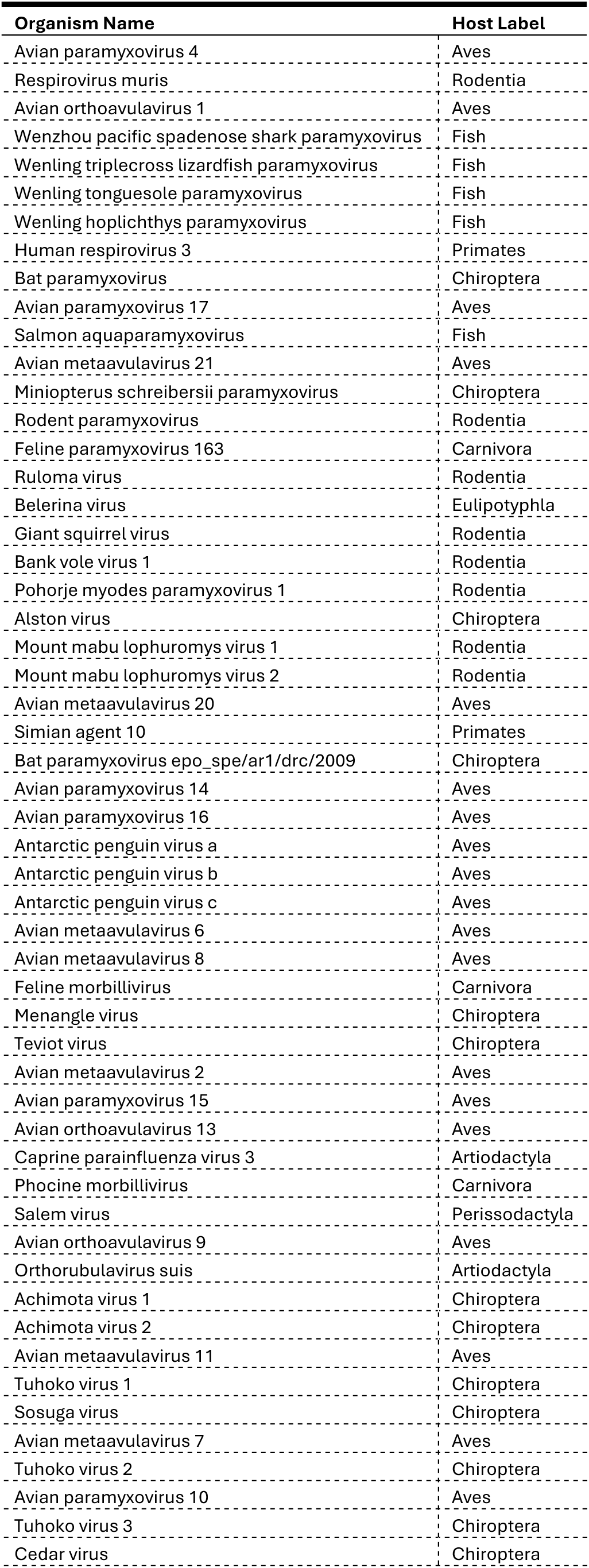
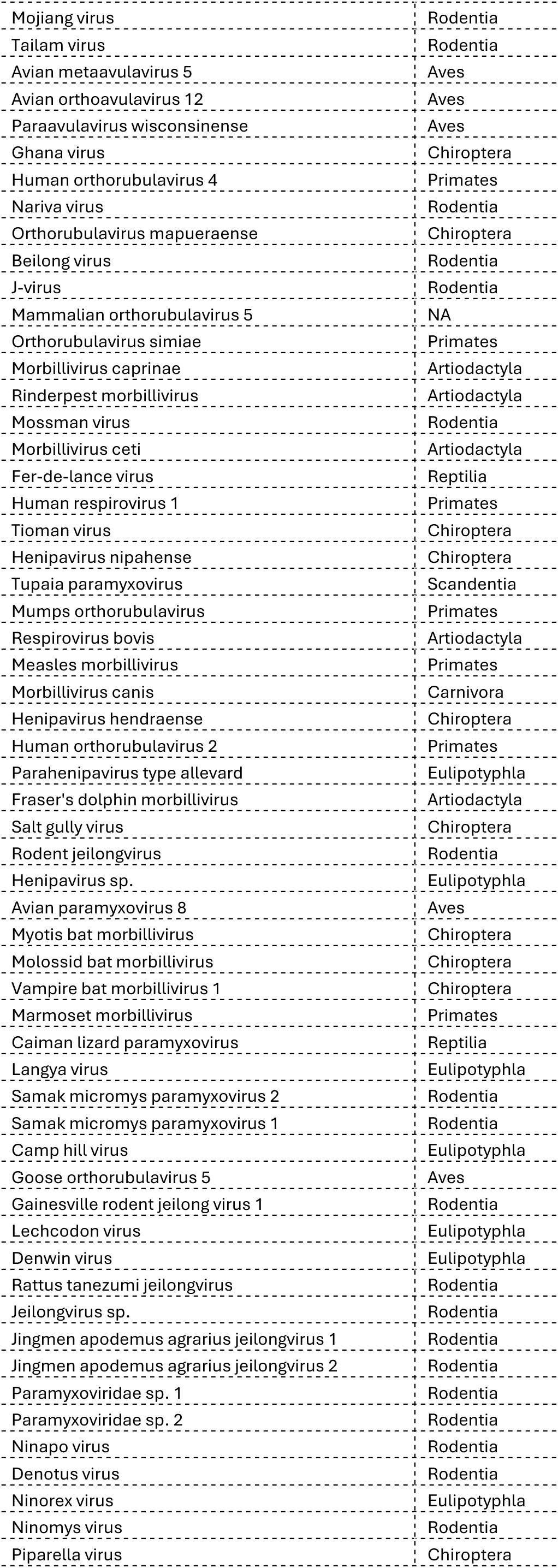
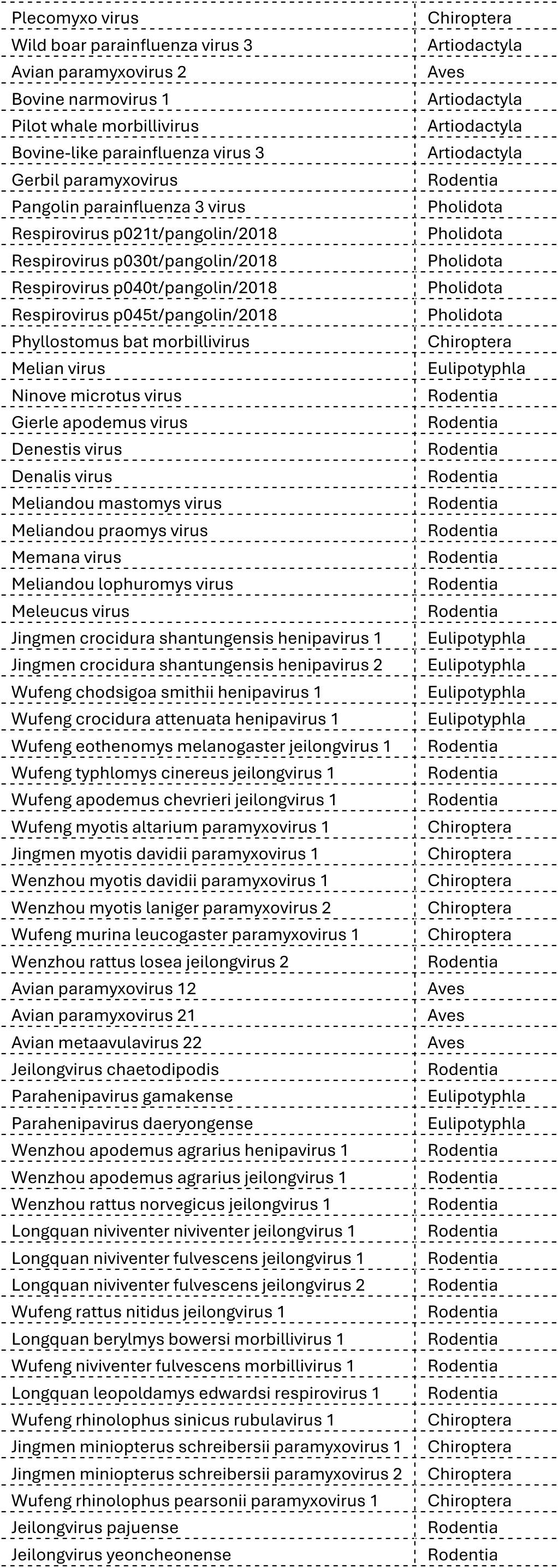
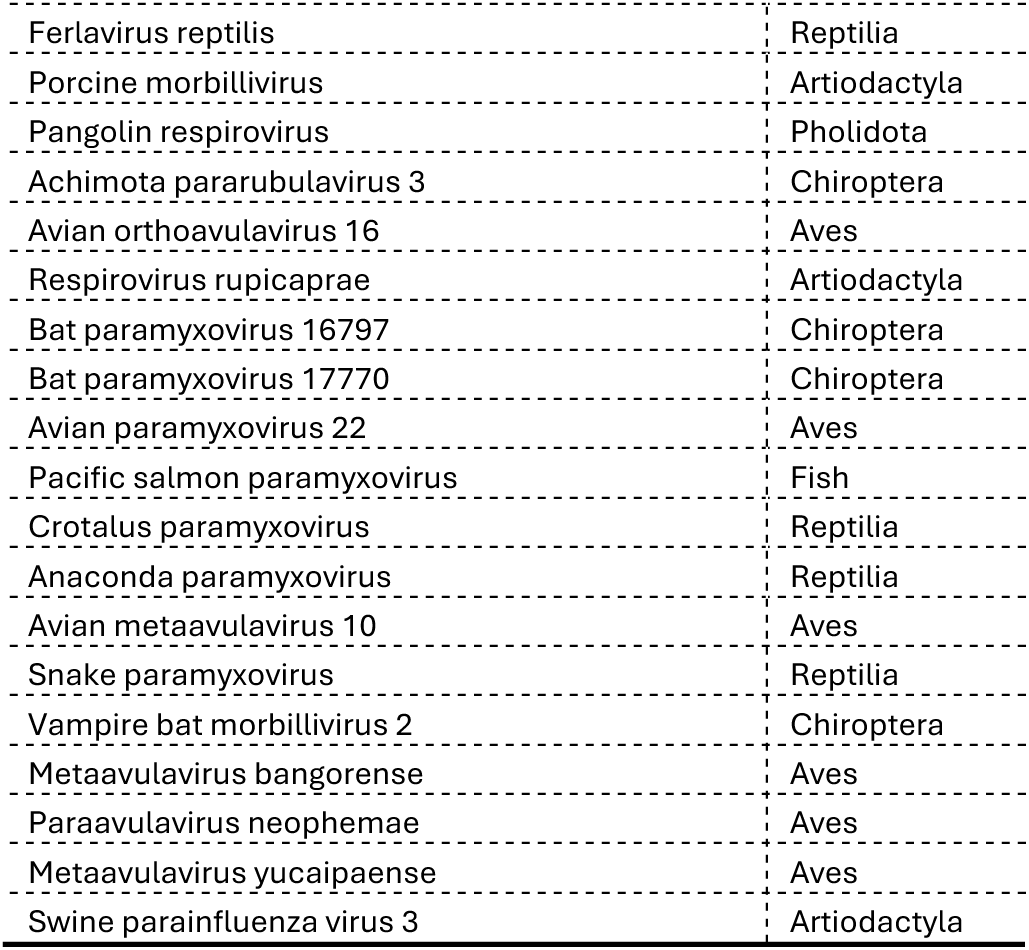
Virus-host lookup table used to assign host labels to viral sequences.

**Supplementary Table 2:** Comparison of pathfinding scores used to determine optimal clock model for BEAST X phylogeny.

|  | Strict | HMC | UCRlog |
| --- | --- | --- | --- |
| **Strict** | 0 | -5,111.84 | -479.02 |
| **HMC** | 5,111.84 | 0 | 4,632.82 |
| **UCRlog** | 479.02 | -4,632.82 | 0 |

## Supplementary Figures

**Supplementary Figure 1:**
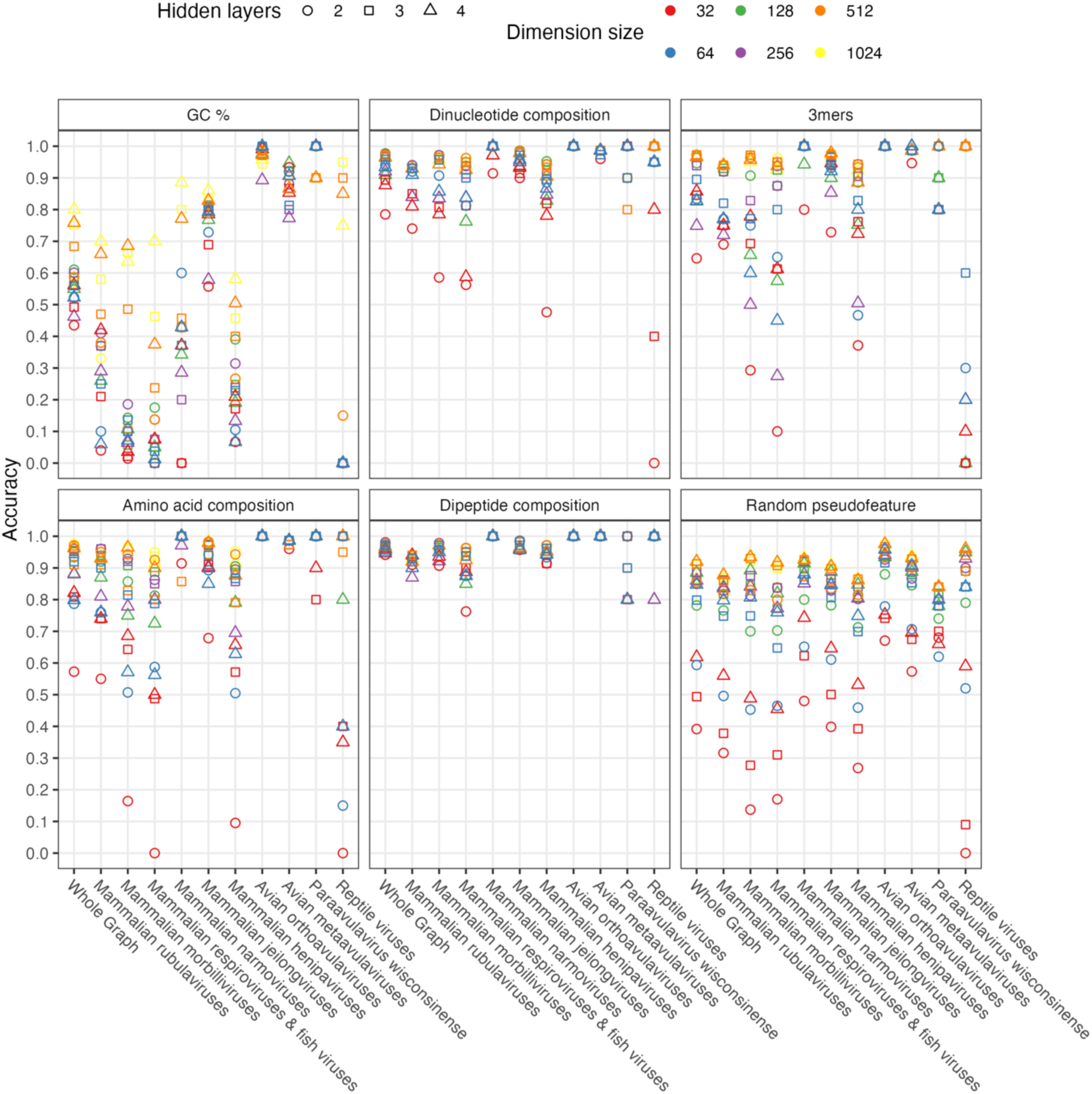
Comparison of GNN models trained using different compositional features across a range of model architectures.

**Supplementary Figure 2:**
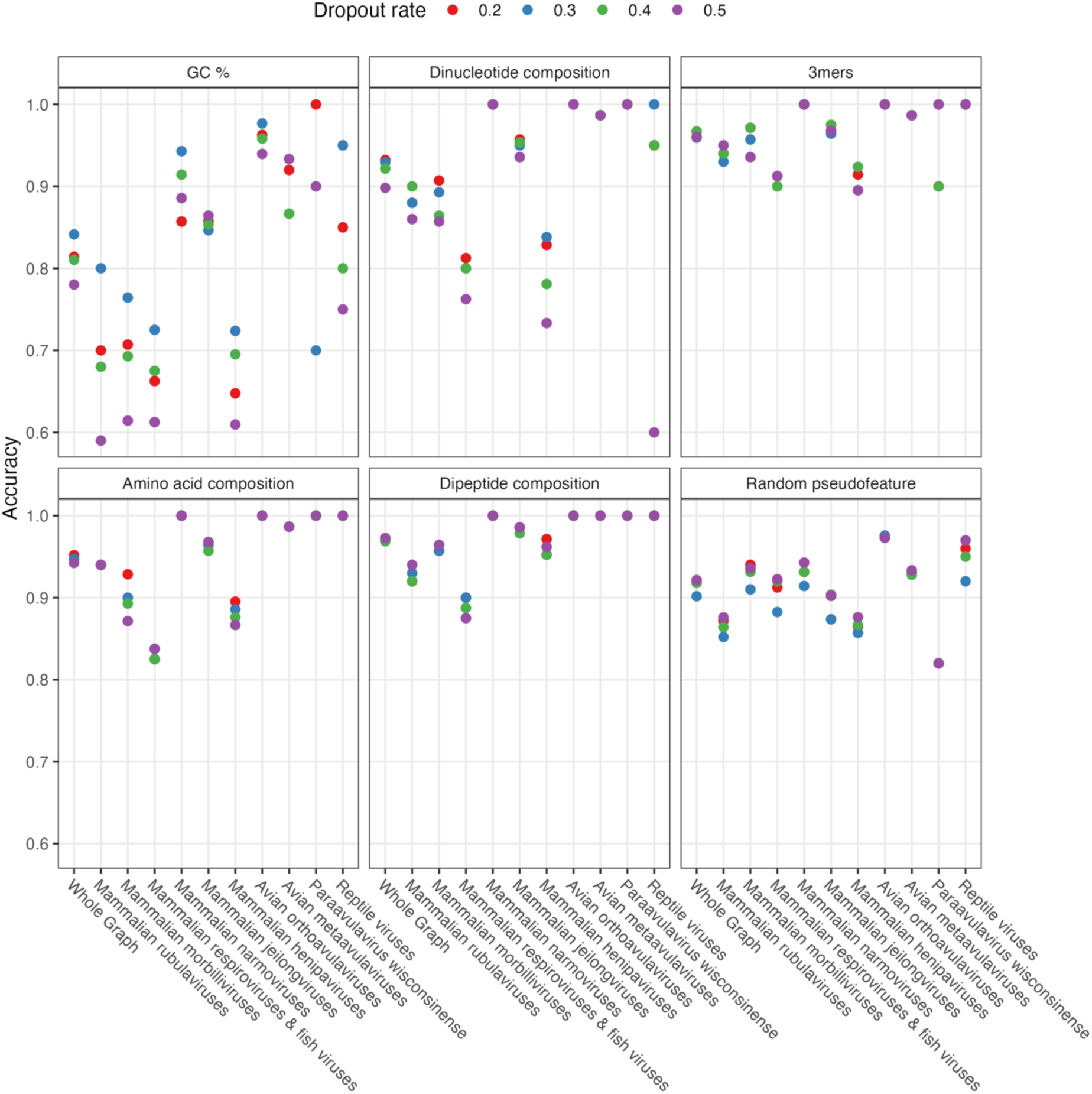
Comparison of performance of GNN models trained using different compositional features with a range of different dropout values.

**Supplementary Figure 3:**
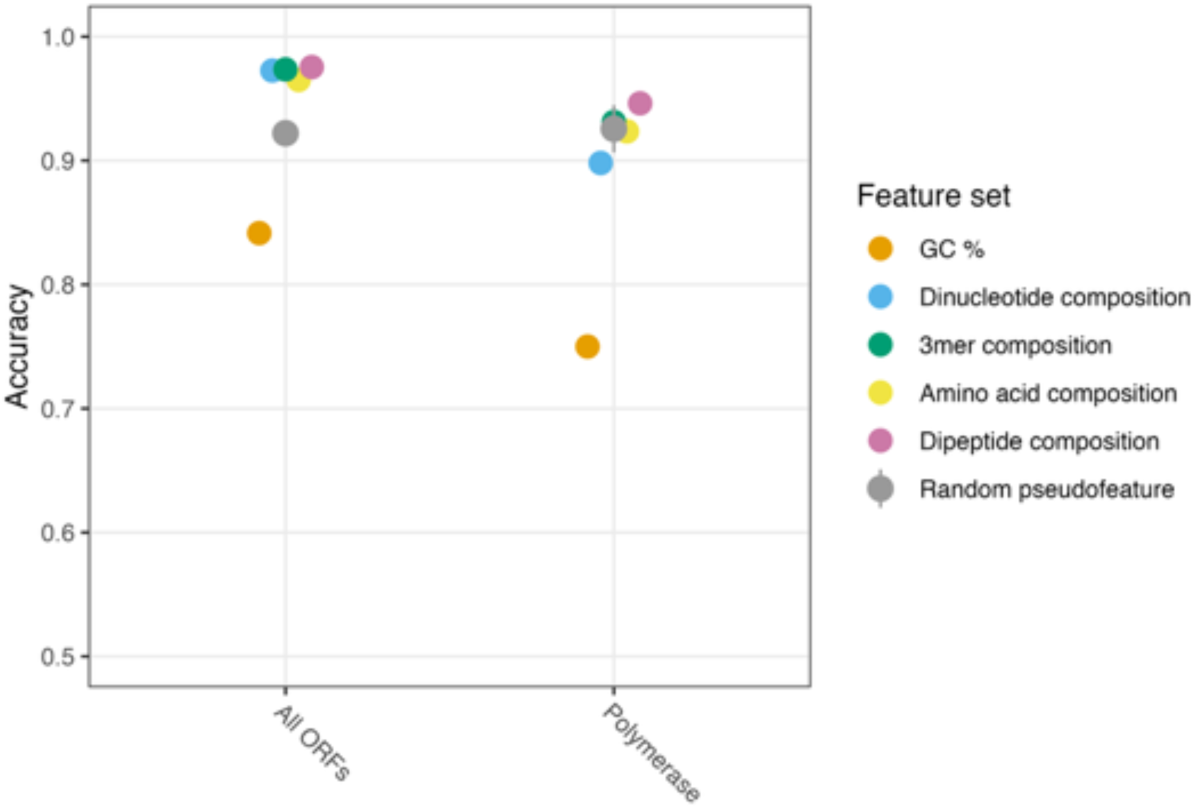
Comparison of performance of GNN models trained using different compositional features with phylogenetic graphs constructed from BEAST phylogenies based on alignments of five canonical ORFs or the L ORF only (coding for the polymerase).

**Supplementary Figure 4:**
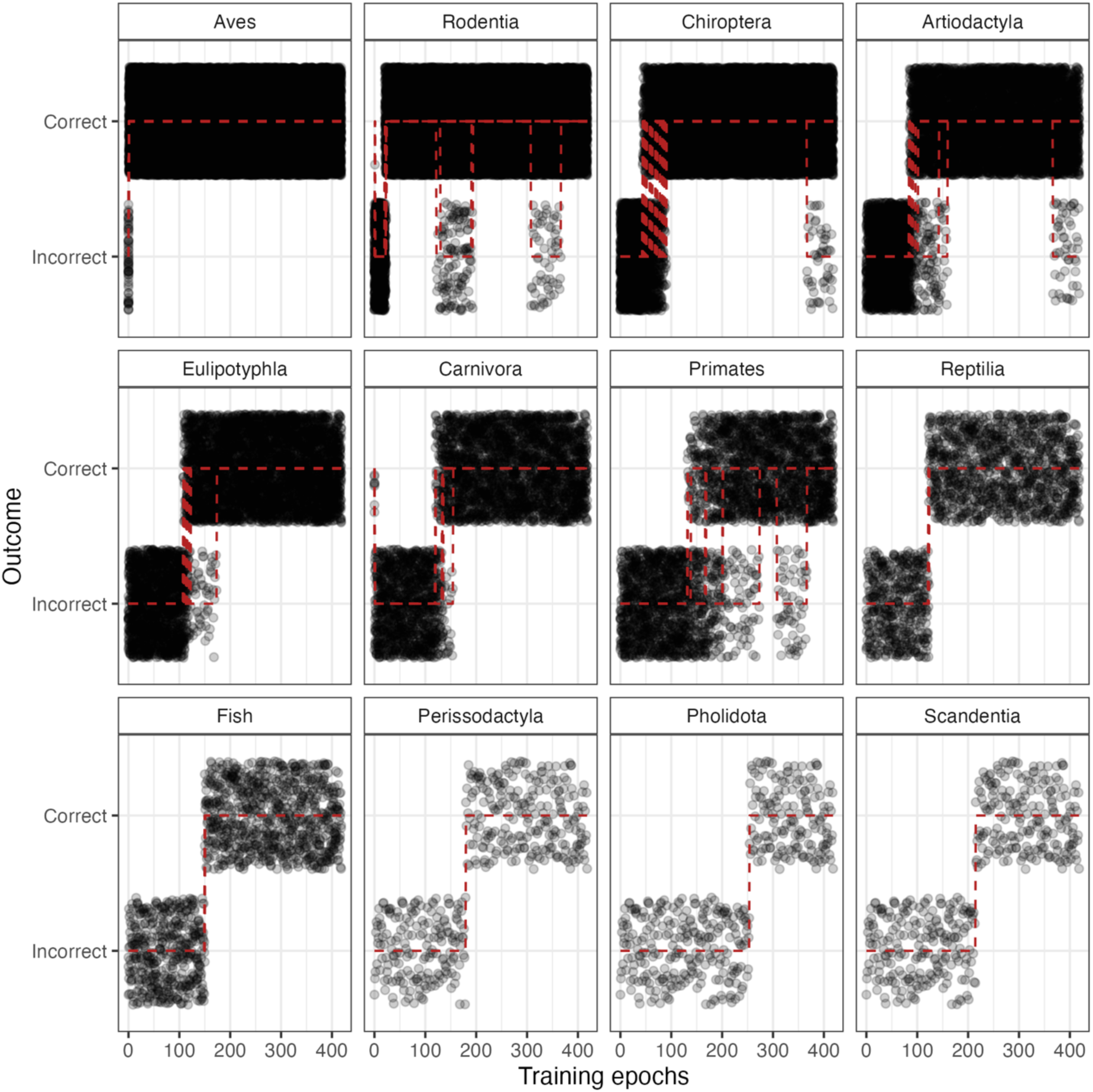
Correct and incorrect predictions of each host label made by GNN model trained using dipeptide compositional features. The prediction on every individual node is plotted as a point at every training epoch. Red dashed lines show individual instances of prediction transitions from incorrect to correct or vice versa. Host labels are presented in order of their frequency in the dataset.

